# CONTROLLING COCKROACH POPULATIONS IN HUMAN AND ANIMAL HABITATIONS BY REDUCING FOOD AVAILABILITY

**DOI:** 10.1101/2021.01.22.427850

**Authors:** William A. Hayward, Luke J. Haseler, Maheswari Muruganandam, James I. Gibb, John H. Sibbitt, Wilmer L. Sibbitt

**Affiliations:** The Department of Exercise and Sport Sciences, New Mexico Highlands University, Las Vegas, NM, USA; Faculty of Health Sciences, School of Physiotherapy and Exercise Science, Curtin University, Perth, Australia. Emil; Department of Internal Medicine, Division of Rheumatology and School of Medicine, MSC 10 5550, 5^th^ FL ACC, University of New Mexico Health Sciences Center, Albuquerque, NM, USA 87131.; Department of Internal Medicine, Division of Nephrology and School of Medicine, MSC 10 5550, 5^th^ FL ACC, University of New Mexico Health Sciences Center, Albuquerque, NM, USA 87131.; Albuquerque, NM, USA 87106.

**Keywords:** cat, pet, cockroach, ant, dish, disease prevention

## Abstract

**Background:** Cockroaches, carriers of pathogenic organisms that cause human and animal disease, typically have access to human or pet food in bowls, allowing cockroaches to expand their colonies and infestations.

**Hypothesis:** We hypothesized that existing anti-ant technology could be converted to anti-cockroach technology by simple design changes.

**Methods:** A base of various heights was affixed to the bottom of an anti-ant bowl to increase the distance of the anti-ant shield from the native height ***“x”*** to the hypothesized cockroach-resistant height ***“z”***. The effects of z =0, 12.7, 15.9, 19.1, 25.4, 44.5, and 57.2 mm were studied. 118.3 cc (4 oz) of dry cat food was used as cockroach bait. The modified anti-ant bowls were placed in a high-intensity cockroach environment during summer nights where the temperatures varied between 23.9-29.4 degrees Celsius for 3 hours and then cockroach counts were performed. Ten runs at each height z were performed.

**Results:** Mean numbers of infesting cockroaches ± SD at each height z were 21.3±2.9 at 0 mm, 22.0±2.9 at 12.7 mm, 11.2±2.6 at 15.9 mm, 0.9±0.8 at 19.1mm, 0.4±0.5 at 25.4 mm, 0±0 at 44.5 mm, and 0±0 at 57.2 mm (p<0.001 with ***z***≥15.9 mm for all). Cockroach numbers began declining when ***z*** = 15.9 mm and declined to only large cockroaches at ***z*** = 25.4 mm. The cockroaches that were able to overcome the ***z*** =25.4 mm were the larger American cockroaches that can exceed 76.2 mm (3 inch) in length. However, at ***z*** = 44.5 mm and 57.2 mm no cockroaches penetrated the modified bowl.

**Conclusions:** To defeat the majority of species of cockroaches the anti-insect shield should be at a height of at least 25.4 mm and to defeat the larger American cockroaches preferably greater than 25.4 mm with 44.5 mm and 57.2 mm defeating all tested cockroaches

## Introduction

Cockroaches commonly infest sewers, buildings, gardens, and any other area where there is warmth and food (1–5). Cockroaches are proven carriers of pathogenic organisms that cause both human and animal disease resulting in vector-borne illness and death. Cockroach carcasses and excrement also cause allergic reactions, including dermatitis, anaphylaxis, and chronic asthma (4). The key to control of cockroaches is to completely block their access to food (5). In this regard, one important source of food for cockroaches in and around human habitations is human or pet food that is commonly left in a bowl on a table, the floor of a kitchen, or on the ground outside providing cockroaches easy access to unlimited food (6). Thus, human or pet food can spawn large colonies of cockroaches both inside and outside of houses and other buildings.

Because ants also commonly infest pet food, there are number of commercial anti-ant pet food bowls that are designed to prevent intrusion by ants (7–9). However, these anti-ant products often fail in preventing intrusion of resident cockroaches into food. This present research addressed the hypothesis that design changes to existing anti-ant food bowls would convert the anti-ant technology of commercial products into anti-cockroach technology to prevent ambient cockroach intrusion.

## Methods

A number of commercial anti-ant food bowls have a protruding anti-ant flange, skirt, or shield above the ground that is situated so that insects, primarily ants, cannot access the outside surface of the shield because the shield exceeds the ant’s reach dimension (7–9). **Figure 1** represents a perspective view of such a typical anti-ant device as described comprising an animal feeding dish including a generally circular shaped bowl assembly having a shield disposed above the ground or floor surface by a base support member. The base support member is generally cylindrical in shape and supports the bowl food holding area above the ground or floor. Generally a rim extends horizontally to anti-ant shield that is a distance “***x***” above the ground so that ants cannot reach or access the external surface of the anti-ant shield and thus cannot access the interior of the bowl.

**Figure 1.**
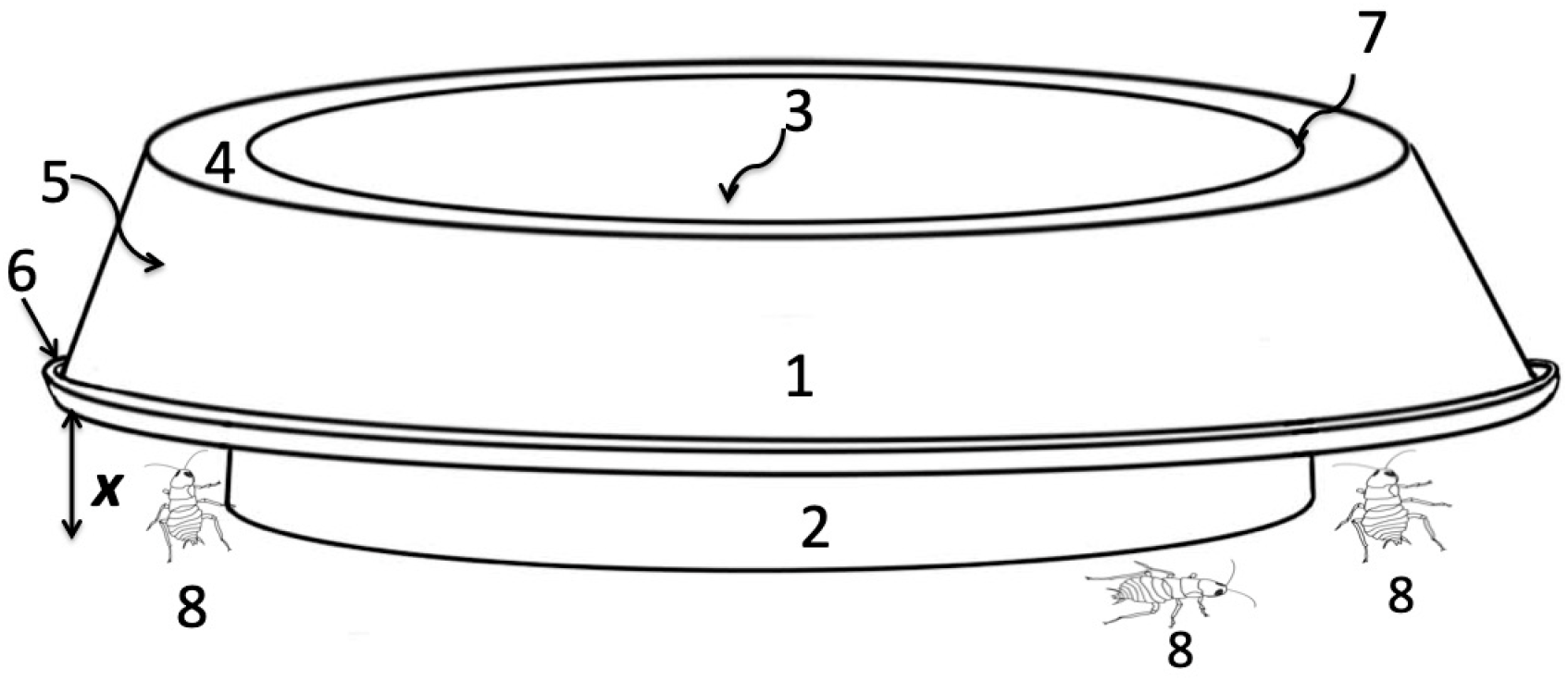
This figure represents a perspective view of such a typical anti-ant device as described by Hand et al (8,9) comprising an animal feeding dish **1** including a generally circular shaped bowl assembly **7** having a disposed above the ground or floor surface by a base support member **2**. The base support member **2** is generally cylindrical in shape and supports the bowl food holding area **3** above the ground or floor. Generally a rim **4** extends horizontally to anti-ant shield or skirt **5** with a lip **6** that is a distance “***x***” above the ground so that ants **8** cannot reach the lip **6** or access the external surface of the anti-ant shield **5** and thus cannot access the extension **4** or the interior of the bowl **3**.

The designs of these commercial anti-ant bowls as in **Figure 1**, however, do not stop flying insects or large insects like certain species of cockroaches that can typically be 25.4 mm (1 inch) up to 76.2 mm (3 inches long) (2,10–12). **Figure 2** shows a common American cockroach that can exceed 80 mm (3.14 inch) in length. Typically, commercial anti-ant bowls do not prevent intrusion of cockroaches. We hypothesized that cockroaches defeat the dimensions of the ant shield by extending from the ground onto the external facing surface of the shield and thus scale into the bowl or scale the wall of the bowl and then reach outward to the edge of the shield and climb into the bowl **(Figure 3).** Thus, we believed that these anti-ant products fail preventing intrusion of cockroaches because the larger size of cockroaches relative to ants allow these larger insects to access the exterior of the anti-ant shield, defeating this barrier, and gaining access to the pet food bowl **(Figure 4).**

**Figure 2.**
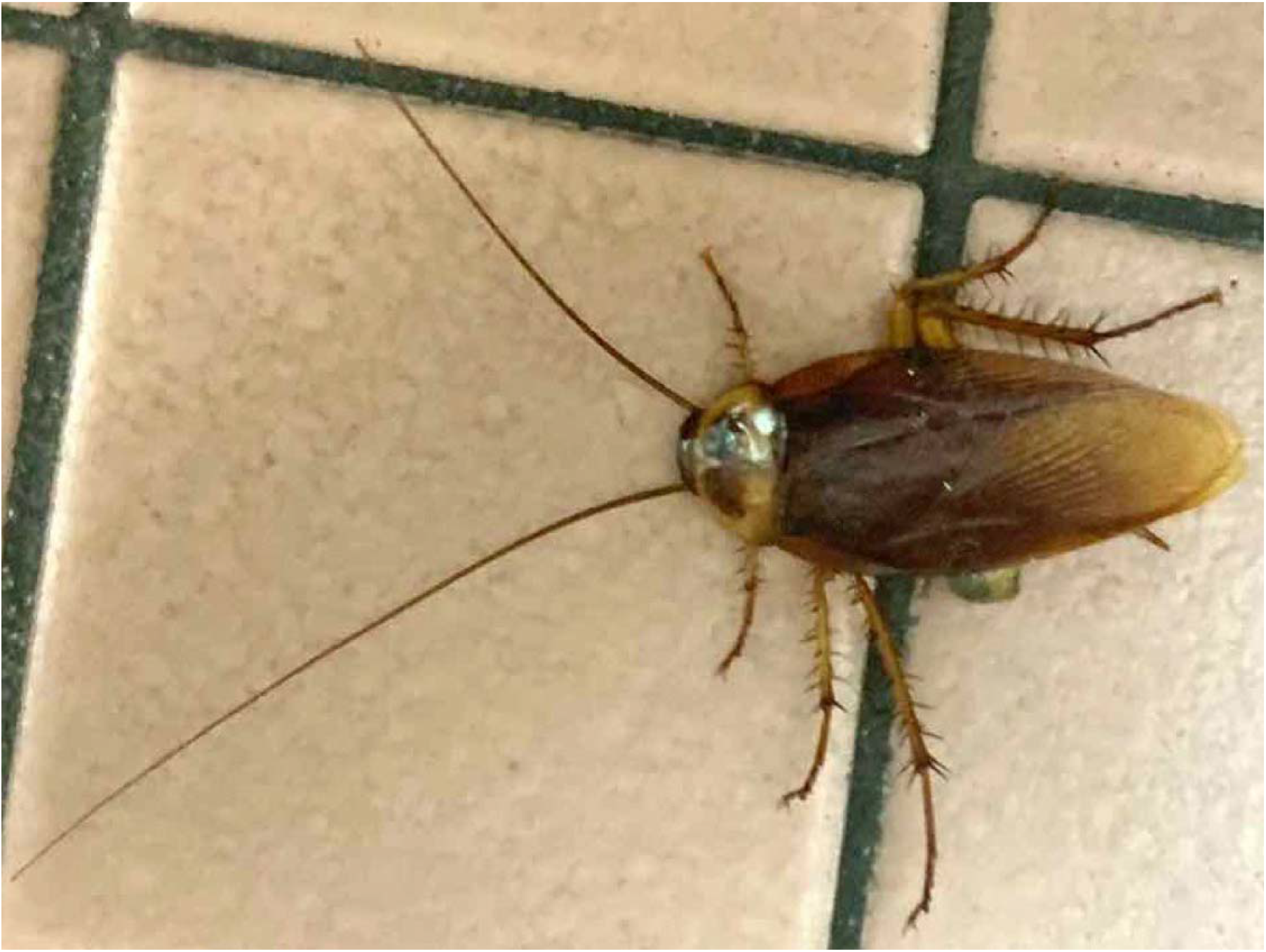
***Periplaneta americana***, the American cockroach, the adult forms of which are typically 35–40 mm (1.4-1.6 inches), but may 80 mm (3.14 inch) in length. The American cockroach is originally from Africa, but is a widespread pest throughout North America and now the world in buildings and sewers.

**Figure 3.**
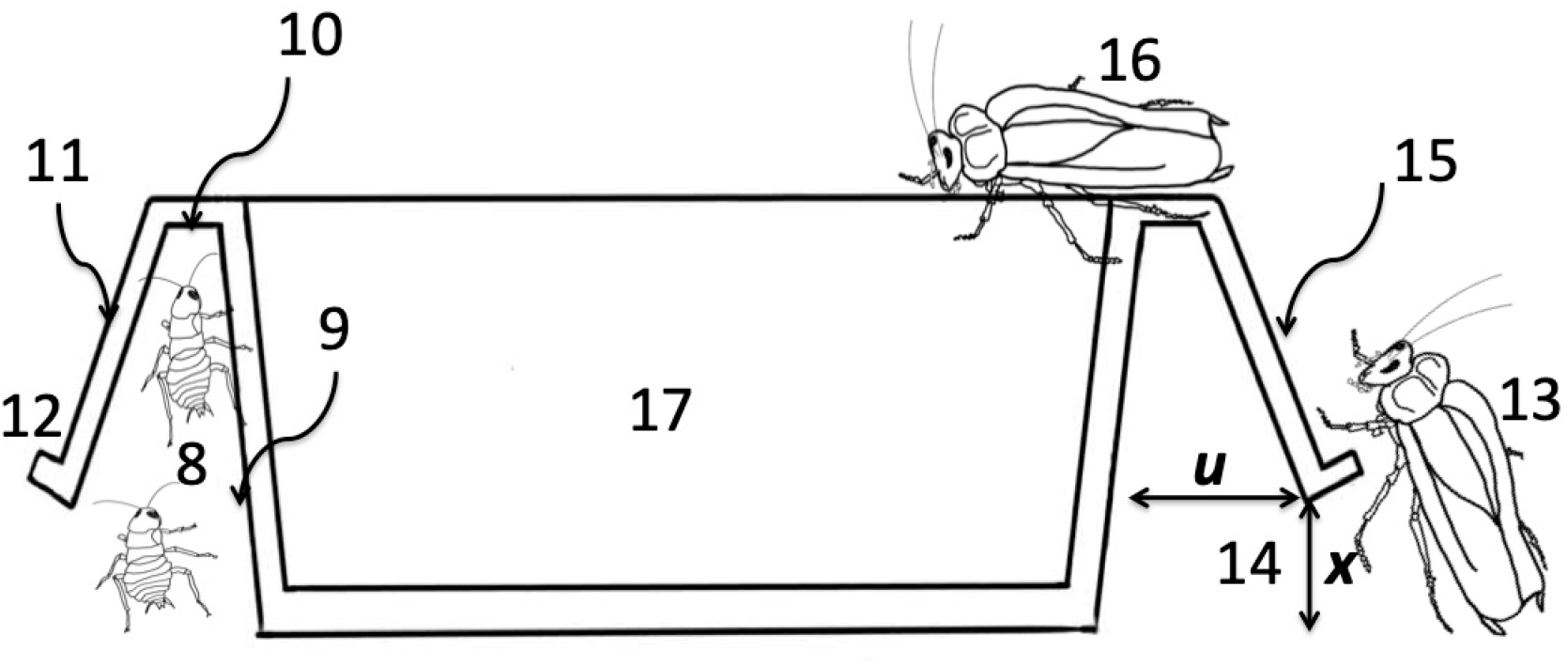
A cross sectional view of a typical commercial anti-ant device as in **Figure 1** ants **8** can access the interior dimension of the shield “***u***” and crawl up the base support **9** but are blocked from entry by the internal surface of rim **10** and cannot access the external surface of the anti-ant shield **11** and lip **12**. However, we hypothesized that large cockroaches **13** who with their legs can exceed the distance “***x***” **14** can access both the external surface of the lip and anti-ant shield, and can scale the external surface of the anti-ant shield **15** and then the cockroach **16** can access the interior of the bowl **17**.

**Figure 4.**
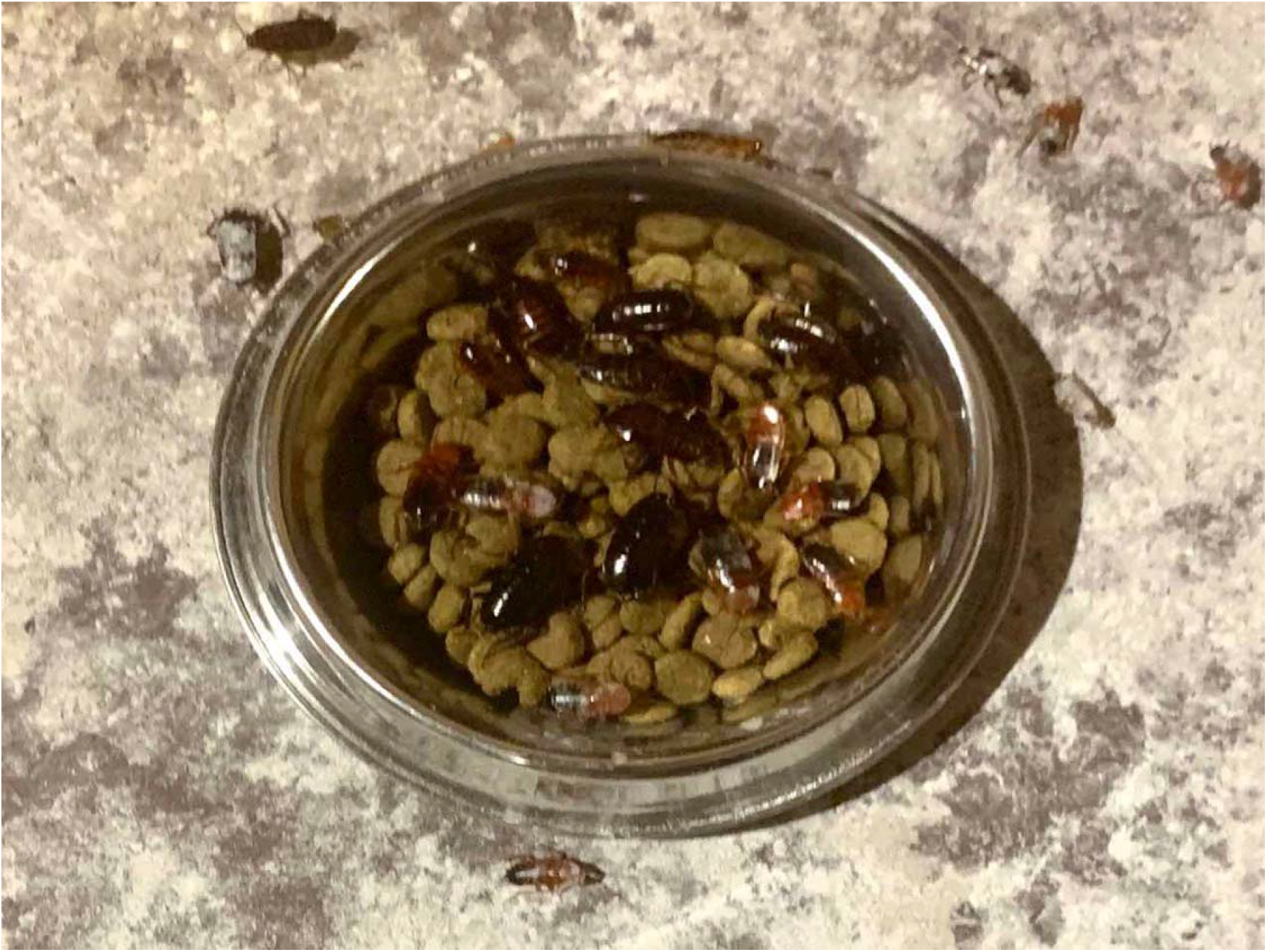
A commercial 236.6 cc (8 oz) anti-ant bowl left in a cockroach-infested area for 3 hours. There are 17 or more cockroaches of various sizes, these being mostly ***Shelfordella lateralis***, the Turkestan cockroaches, that have replaced ***Blattella germanica***, the German cockroach, in many locales.

A 236.6 cc (8 oz) anti-ant bowl (Anti-Ant Stainless Steel Non Skid Pet Bowl for Dog or Cat –8 oz –1 cup, SKU 799665921910, Item Number 92191, Iconic Pet, LLC, 611 South Ave, Garwood, NJ 07027. Website: www.iconicpet.com) was studied to determine how an anti-ant bowl could be modified to become an anti-cockroach bowl. We hypothesized that an anti-ant bowl could be modified to become an anti-cockroach bowl by affixing a base of a height of ***y*** to the circular plate bottom surface and thus increase the distance of the lip and shield from the native anti-ant height ***x*** to the cockroach resistant height ***z*** that would exceed the ability of the cockroach to access the outside surface of the shield **(Figure 5).** We added and subtracted the base through graduations so that the effects of *z* = 0, 12.7, 15.85, 19.05, 25.4, 44.45, and 57.15 mm (0, 0.5, 0.624, 0.75, 1.0, 1.75, and 2.25 inches) were studied. In these experiments 118.3 cc (4 oz) of dry cat food (Crave with Protein from Salmon & Ocean Fish Adult Grain-Free Dry Cat Food, Crave Pet Foods, Mars Petcare US Company, Franklin, Tennessee, USA, Website: www.cravepetfoods.com) was used as cockroach bait. The modified dishes were placed in a high-intensity cockroach environment during the night during summer where the temperatures varied between 23.9-29.4 degrees Celsius (75-87 degrees Fahrenheit) for 3 hours and then cockroach counts were performed **(Figure 4).** Ten runs at each height ***z*** were performed. After each experiment the cockroaches were not destroyed, but rather released back alive into the high intensity cockroach environment.

**Figure 5.**
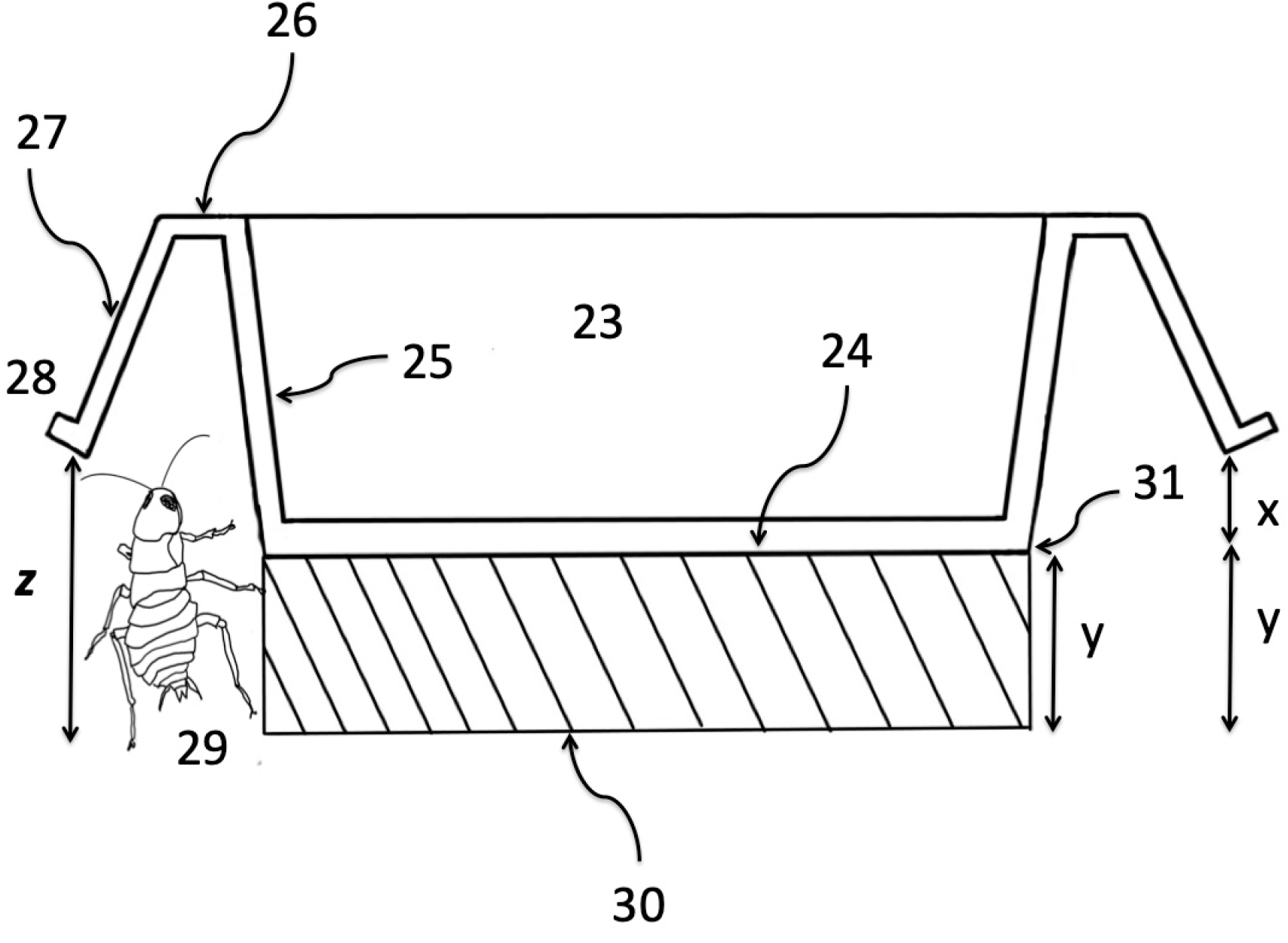
This figure represents an anti-ant bowl as in **Figures 1,3 and 4** that is modified to become an anti-cockroach bowl. The device consists of an internal container area **23**, a circular plate bottom surface **24**, a generally cylindrical sidewall base support **25**, a rim **26**, an anti-ant shield **27**, and a lip **28**. The surface of the lip **28** and shield **27** need to be above the ground a distance “***z***” to exceed the reach of the cockroach **29**. The dimension “***z***” depends on the size and athleticism of the cockroach **29**. A base **30** of a height of “***y***” is affixed to the circular plate bottom surface **24** and increases the distance of the lip **28** and shield **27** from the native height “***x***” to the cockroach resistant height “***z***” that would exceed the ability of the cockroach to access the outside surface of **28** and shield **27**.

### Statistical Methods

Standard summary statistics (means, proportions) were calculated and statistical analysis was performed with Simple Interactive Statistical Analysis (SISA) (Consultancy for Research and Statistics, Lieven de Keylaan 7, 1222 LC Hilversum, The Netherlands; http://www.quantitativeskills.com/sisa/). Summary data were expressed as mean ± standard deviation (SD) and means at different heights *z* were compared with the student t-test with corrections for multiple comparisons. Statistical differences between measurement data were determined with Student t-test. P values <0.05 were considered significant.

## Results

**Table 1 and Figure 6** demonstrate the mean number of cockroaches after a 3-hour period that were able to overcome the anti-ant shield and infest the bowl versus the distance ***z*** as defined in **Figure 5.** Mean numbers of infesting cockroaches ± SD at each height z were 21.3±2.9 at 0 mm, 22.0±2.9 at 12.7 mm, 11.2±2.6 at 15.9 mm, 0.9±0.8 at 19.1mm, 0.4±0.5 at 25.4 mm, 0±0 at 44.5 mm, and 0±0 at 57.2 mm (p<0.001 with ***z***≥15.9 mm for all) **(Table 1).** As can be seen, the number of cockroaches began declining when ***z*** = 15.9 mm and declined to only a few very large cockroaches at ***z*** = 25.4 mm. The cockroaches that were able to overcome the ***z*** =25.4 mm were the larger American cockroaches that can be up to 76.2 mm (3 inches) in length. However, at ***z*** = 44.5 mm and 57.2 mm no cockroaches penetrated the modified bowl **(Figure 7).**

**Figure 6.**
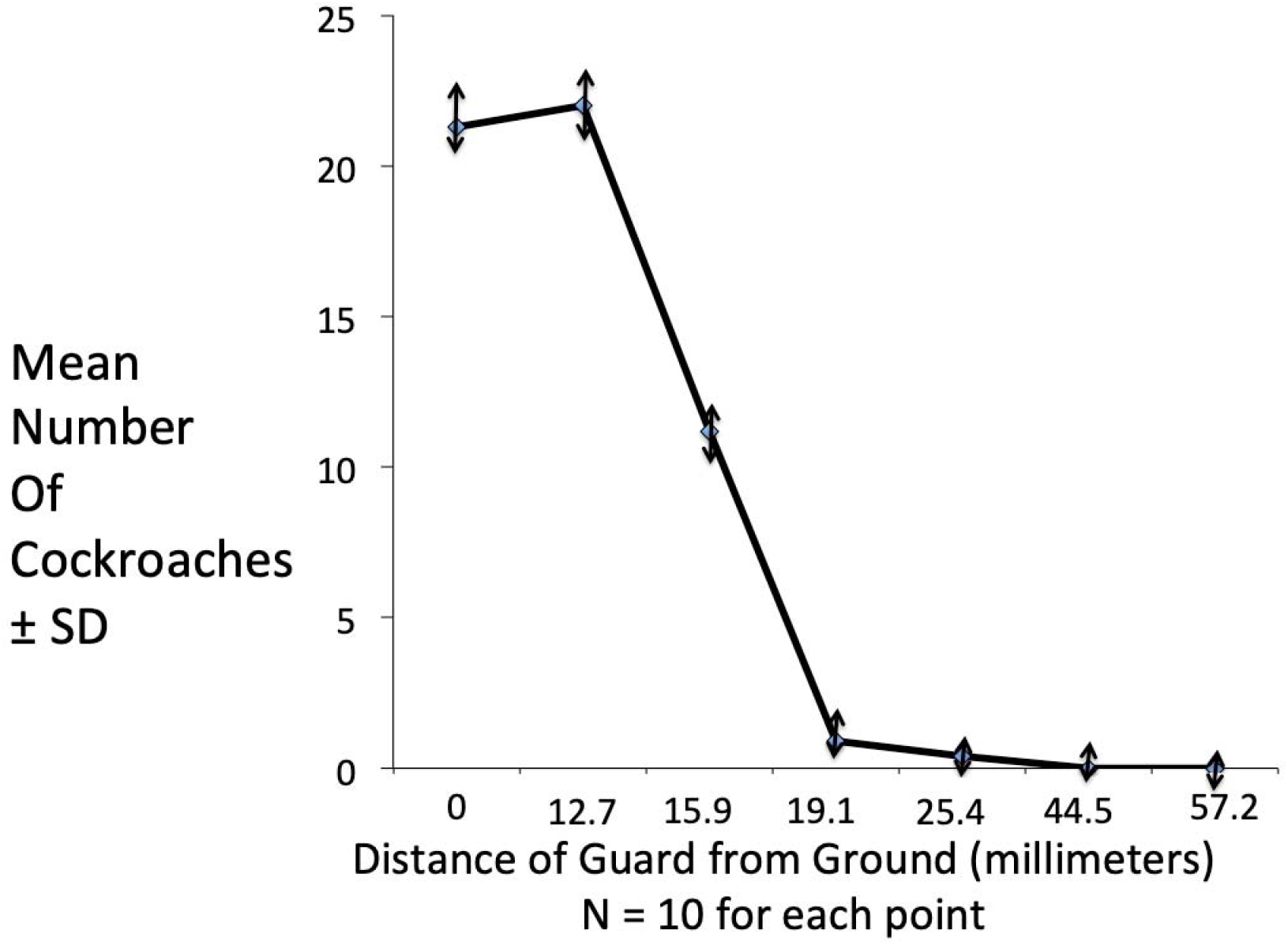
demonstrates the mean number of cockroaches after a 3-hour period that were able to overcome the anti-ant bowl versus the distance “***z***”. As can be seen, the number of cockroaches declined when ***z*** = 15.9 mm (0.625 inch) and declined to only a few very large cockroaches at ***z*** = 25.4 mm (1.0 inch). The cockroaches that were able to overcome the ***z*** =25.4 mm (1.0 inch) were the larger American cockroaches that can be greater than 1 inch up to 3 inches in length. However, at ***z*** = 44.5 mm (1.75 inches) and 57.2 mm (2.25 inches) no cockroaches penetrated the modified bowl.

**Figure 7.**
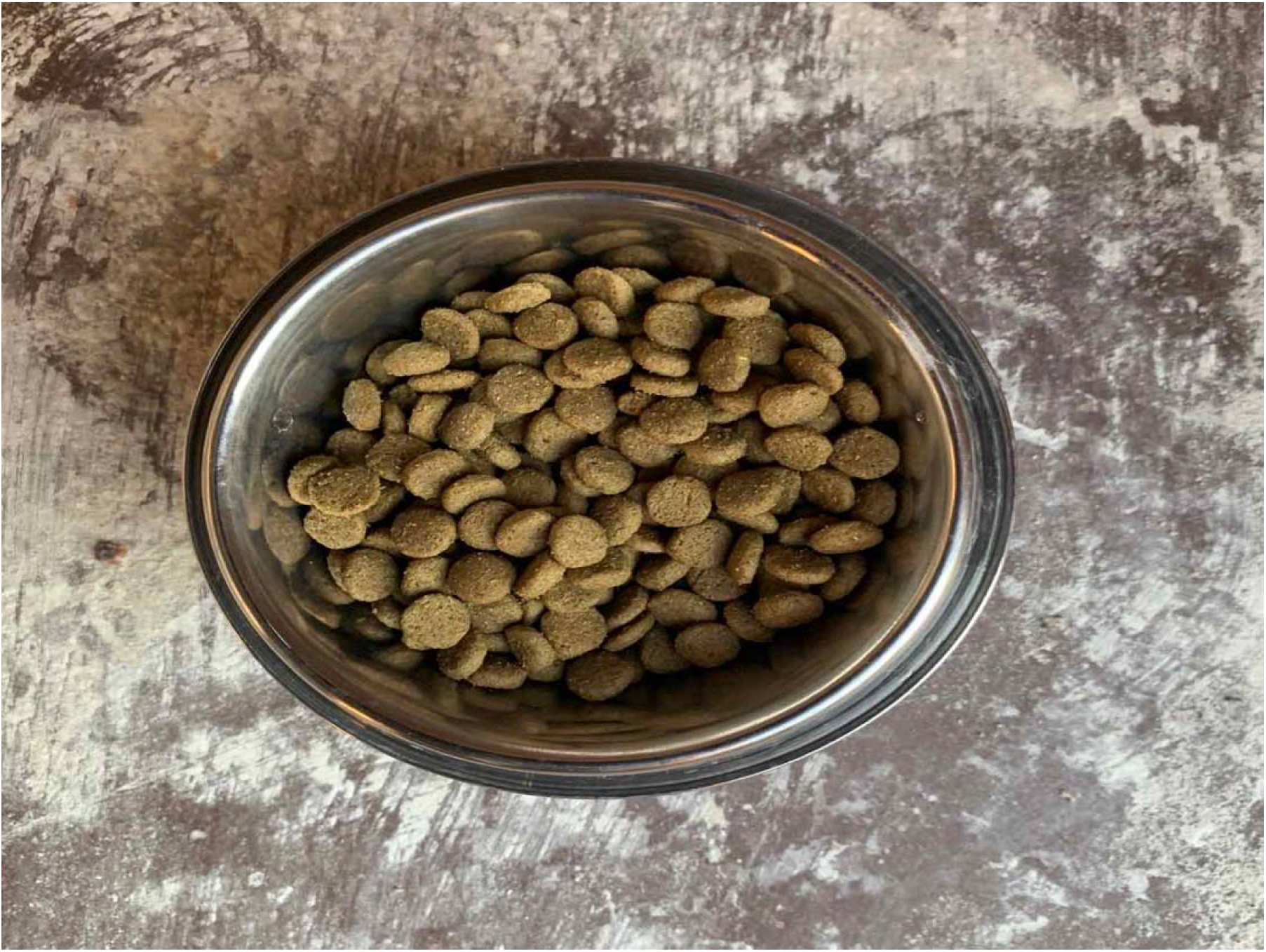
A 236.6 cc (8 ozĵ anti-ant bowl with a modified base so that ***z*** = 57.2 mm (2.25 inches) left in a cockroach-infested area for 3 hours. There is no cockroach intrusion.

**Table 1.** Number of Cockroaches versus Height of Anti-Ant Shield

|  |  |  |  |  |  |  |  |
| --- | --- | --- | --- | --- | --- | --- | --- |
| Height of shield (mm) | 0 | 12.7 | 15.9 | 19.1 | 25.4 | 44.5 | 57.2 |
| Number of runs | 10 | 10 | 10 | 10 | 10 | 10 | 10 |
| Meant number of cockroaches | 21.3 | 22 | 11.2 | 0.9 | 0.4 | 0 | 0 |
| Standard Deviation | 2.9 | 2.9 | 2.6 | 0.77 | 0.5 | 0 | 0 |
| 95% CI of difference (Wald): | 3.4<0.7<2.0 | NA | 13<11<8 | 23<21<19 | 24<22<19 | 24<22<20 | 24<22<20 |
| *P value | 0.6 | NA | <0.001 | <0.001 | <0.001 | <0.001 | <0.001 |

CI = confidence interval
\*P values were determined with the t-test using the cockroach numbers at the native height of the bowl at 12.7 mm as a comparator. Corrections were made for multiple comparisons.

Thus, to defeat the majority of species of cockroaches the anti-ant shield should be at a height of at least 25.4 mm and to defeat the larger American cockroaches preferably greater than 25.4 mm with 44.5 mm and 57.2 mm defeating all tested cockroaches

## Discussion

Because cockroaches feed on human and animal feces these insects commonly spread bacteria, viruses, and parasites known to cause both animal and human disease. Cockroaches are proven carriers of pathogenic organisms including staphylococcus, enteric organisms, streptococcus, viruses and parasites – organisms that may cause life-threatening diarrhea, dysentery, cholera, leprosy, plague, typhoid fever and viral diseases such as poliomyelitis resulting in severe illness or death (1–5). In addition cockroaches carry the invertebrate parasites, including eggs and cysts of parasitic worms and organisms that infest both humans and pets. Cockroach carcasses and excrement may also cause severe allergic reactions, including dermatitis, itching, swelling of the eyelids, anaphylaxis, and allergic asthma, resulting in increased costs, medical care, and in some cases death (4). Thus, an inexpensive intervention that reduces the cockroach burden in areas of high human population has the potential for major public health benefit. The results reported here are in support of our hypothesis and confirm that a properly designed anti-insect shield can prevent cockroach access to pet food as a nutrient source.

Cockroaches are insects from the size of large ants 2–3 mm (0.08- to 0.12 inch) to the size of large beetles over 80 mm (3.14 inch) in length (1–6) **(Figures 2 and 4).** Of over 3500 identified cockroach species only a few have adapted to living in buildings in close association with people and these cockroaches have become serious pests (1–6). Cockroaches eat crumbs, pet food, cookies on a plate, human and animal feces and even human skin and nail clippings (2,6).

The key to control of cockroaches is to remove their access to food (2). In this regard, one very important source of food for cockroaches in and around human habitations is human or pet food, especially dog and cat food, that is commonly left in a bowl on the ground outside or on floor of a kitchen for the pets where the cockroaches can and do easily access the pet food (6) **(Figure 4).** Thus, pet food can spawn large colonies of cockroaches both inside and outside of houses. The cockroaches contaminate the pet food with their feces and secretions and thus transmit bacterial, parasitic, and viral diseases to both the pets and to the pet’s human owners. Because of the excess of food, the cockroaches are able to expand their colonies and cockroach numbers, causing an even greater infestation locally and the cockroach colonies may expand both inside and outside the house and then invade adjoining properties.

Because of the magnitude of the problem of crawling insects, in particular ants, infesting dog and cat food bowls, there are many patents and products relating to insect-proof pet bowls (7,13–19). A number of different mechanisms are used to prevent intrusion of insects, particularly ants, into the food bowl. One solution to preventing ingress of ants has been the use of a barrier consisting of a moat that is filled with water, insecticide, or other form of insect repellant (13–15). A problem with these moats or traps is that they must be filled regularly with water or liquid to function – once they dry out, they no longer repel insects. Further, the moats typically catch and drown many of the crawling insects, so eventually the traps fill with the rotting carcasses of insects that must be periodically cleaned from the moats. Further, if the moat is also used as a drinking station the drinking water becomes contaminated with all the filth, bacteria, viruses, and parasite eggs that insects and cockroaches carry. Further, certain cockroaches easily survive in watery environments like sewers and thus are semi-aquatic and can easily defeat liquid barriers.

Another solution to prevent ingress of insects is the use of noxious substances such as caustic chemicals, insecticides, and insect repellants (16–18). A problem of using noxious substances is that the pet or human may be injured, killed, or made sick by the caustic chemical, insecticides, and repellants. Further, the noxious substance must be regularly replenished.

Another solution is a mechanical barrier on the dish or bowl that prevents ingress of crawling insects consisting of a protruding anti-ant flange, skirt, or shield that is situated a certain distance above the ground so that ants cannot access the outside surface of the shield and thus cannot enter the bowl **(Figure 1)** (8,9,19). This basic design is presently used in many contemporary anti-ant pet feeders. This design, however, does not stop flying insects or, as the present research demonstrates, large insects like certain species of cockroaches that can defeat the dimensions of the ant shield by extending from the ground onto the external facing surface of the shield and thus scale into the bowl or scale the wall of the bowl and then reach outward to the edge of the shield and climb into the bowl **(Figures 2 and 3).** Typical commercial “ant-free” products based on this design that were tested in the present research did not prevent cockroaches from accessing the interior of the bowl **(Figure 4).** The results of the present research demonstrate that many cockroaches can still access the bowl at 19 mm (0.75 inch) and greater off of the ground **(Figures 4 and 6)** (8,9). We hypothesized that the failure of these anti-insect designs that are effective for ants, but fail for cockroaches is largely due to the larger dimensions and athleticism of cockroaches that defeat these mechanical barriers **(Figures 2,3,4,6, and 7)**.

The most common cockroach species considered pests in human habitations are as follows:

***Periplaneta americana***, the American cockroach, the adult forms of which is 35– 40 mm (1.4-1.6 inches), but may exceed 51 mm (2 inches) in length up to 80 mm (3.14 inch) in length **(Figure 2).** The American cockroach is originally from Africa, but is a widespread pest throughout North America and the world in buildings and sewers.
***Periplaneta australasiae***, the Australian cockroach, which is similar to the American cockroach and is 31–37 mm (1.2-1.5 inch) long.
***Blatta orientalis***, the Oriental or Chinese cockroach, found mainly in cool temperate regions. It is blackish and 20–27 mm (0.8-1.1 inch) long.
***Supella longipalpa***, the brown-banded cockroach, 10–14 (0.4-0.6 inch) mm long and has yellow and brown bands
***Blattella germanica***, the German cockroach, found in most parts of the world. It is light yellowish brown and 10–15 mm (0.4-0.6 inch) in length.
***Shelfordella lateralis (Blattella lateralis)***, the Turkestan cockroach the females are 20–36 mm (0.8-1.4 inch) in length. It is light yellowish brown. The males are 10–15 mm (0.4-0.6 inch) in length (1–6, 10–12, 20–22).

As can be seen from the above, 4 of the 6 common pest cockroach species (the American cockroach, the Australian cockroach, the Oriental cockroach, and the Turkestan cockroach) commonly exceed 19 mm (0.75 inch). Indeed in the present research anti-ant products based on these designs failed when tested against cockroaches **(Figures 4 and 6, Table 1).** Thus, to be able to defeat the American, Australian, Oriental, and Turkestan cockroaches the barrier must be able to exclude cockroaches much larger than 19 mm (0.75 inch).

We hypothesized we could modify an anti-ant bowl to become an anti-cockroach bowl by affixing a base to the bowl and thus increase the distance of the anti-ant shield from the native anti-ant height to a cockroach-resistant height that would exceed the ability of the cockroach to access the outside surface of the shield **(Figure 5).** As can be seen in **Figure 6**, the number of cockroaches intruding began declining when the height of the anti-ant shield was 15.9 mm (0.625 inch) and declined to only a few very large cockroaches when the height of the shield was 25.4 mm (1 inch). The cockroaches that were able to overcome 25.4 mm (1 inch) were mostly the larger American cockroaches that can be up to 76.2 mm (3 inches) in length **(Figure 2)** (10–12). However, at a height of 44.5 mm (1.75 inches) and 57.2 mm (2.25 inches) no cockroaches penetrated the modified bowl **(Figures 6 and 7).** Thus, to defeat the majority of small to medium cockroaches should the shield should be at least 25.4 mm (1 inch) or greater and to defeat the larger American cockroaches preferably greater than 25.4 mm (1 inch) with 44.5 mm (1.75 inches) and 57.2 mm (2.25 inches) defeating all tested cockroaches **(Figures 6 and 7, Table 1).** Further, the anti-cockroach modifications do not interfere with the anti-ant properties of the bowl.

The reason the tested anti-ant and anti-cockroach shield so effectively interferes with the movement of insects into the bowl is uncertain. Clearly all of the insects have the physical ability to walk on surfaces of the bowl upside down where they could climb up the side of the bowl, walk upside down on the inferior surface of the shield away from the center of the bowl, wrap themselves around the lip, and then access the superior surface of the shield and walk into the bowl **(Figures 1 and 3).** However, the insects tested in the current experiments did not do this. Hand et al speculated that the orientation of an insect shield creates a mechanical barrier that disorients the insect’s foraging activity, increases the insect area restricted search time making defeating the shield unacceptably time-consuming, disrupts communication between insects and thus cooperative foraging, interferes with trail pheromones of insects that successfully reached the bowl area, and attenuates the polarized and unpolarized ultraviolet light used for navigation and orientation by insects thus defeating their access to the bowl (8,9). However, the present research demonstrates the anti-ant shield is also highly effective as an anti-cockroach shield when properly elevated to account for the larger cockroach size **(Figures 4–7, Table 1).**

The geographical ranges of pest cockroaches include most of the world and are especially concentrated in urban areas (1–6). Pest cockroaches can be found in tropical Lagos, Nigeria as well as in Moscow, Russia both typically infesting buildings and sewer systems (21, 22). This simple inexpensive mechanical anti-cockroach technology should help prevent pet feeders and human dishes from being a food source for cockroach colonies while still maintaining anti-ant properties and thus decrease infestations and potential transmission of pathogenic organisms to both pets and their human owners.

